# Renal-Derived Human sPRR Does Not Increase Blood Pressure in High Fat Diet Mice

**DOI:** 10.1101/2024.01.30.577981

**Authors:** Gertrude Arthur, Katherine Biel, Jeffrey L Osborn, Terry D. Hinds, Ming Gong, Analia S. Loria

## Abstract

Obesity is a risk factor for hypertension. Obesity-related hypertension has been associated with elevated plasma soluble prorenin receptor (sPRR) particularly in men. Additionally, renal PRR and sPRR protein expression is upregulated during obesity and diabetes. However, whether renal-derived human sPRR (HsPRR) may influence the intrarenal RAS status to regulate blood pressure and kidney function during obesity has not been investigated. Therefore, we studied the role of collecting duct (CD) derived-HsPRR on blood pressure and kidney function in male and female mice during obesity. Eight-week-old male and female CD-HsPRR mice were placed on a high fat diet for 25 weeks. HsPRR increased renal sPRR concentration but did not change its circulating levels in male and female littermates compared to CTL mice. GFR, water intake and urine flow were not influenced by the CD-HsPRR expression in either sex. Moreover, after 21 weeks of HFD, blood pressure was similar between groups, while only male CD-HsPRR mice showed an impaired pressor response to losartan. In the renal cortex, male CD-HsPRR mice showed increased renin and AT1R mRNA expression associated with increased AQP2, and ENaC subunits protein expression. These data indicate that renal-derived HsPRR induces local upregulation in renin, AT1R and sodium/water transporters in male mice without altering renal hemodynamics or blood pressure. In obese females, CD-HsPRR expression did not affect blood pressure or renal function, which suggests that females may be protected from obesity induced renal function impairment and hypertension.

## Introduction

The prevalence of obesity is increasing worldwide. According to the World Health Organization (WHO), about 39% of the world’s populace is overweight, of which more than 13% are obese (1). In the United States, 2 in 5 adults are obese, accounting for 41.9% of the population (2). Obesity is associated with a threefold higher prevalence of hypertension and is a risk factor for high blood pressure and cardiovascular diseases (3).

Obesity-related hypertension is multifaceted in pathophysiology. Obesity leads to insulin resistance and hyperinsulinemia, leptin resistance and hyperleptinemia, sympathetic nervous system (SNS) activation and activation of the renin angiotensin system (RAS), as well as structural and functional changes to the kidney (4–6). These pathways work both individually and in concert to increase blood pressure and exacerbate hypertension (4, 5).

The RAS plays a critical role in the regulation of blood pressure, and sodium and water homeostasis (7, 8). Over the past decade, the prorenin receptor (PRR) has emerged as a new component of the RAS (9) PRR is a complex and multi-functional 350-amino acid protein that exists in three different forms: the full-length transmembrane protein, the soluble PRR (sPRR), and the truncated form (10) The full-length PRR can bind to renin and prorenin, and participate to the generation of Angiotensin (Ang)-I by increasing the catalytic activity of renin and promoting prorenin nonproteolytic activation (11, 12) PRR can also initiate intracellular signals, especially the activation of MAP kinases through an Ang-II-independent pathway (13),(14) Furthermore, PRR can interact with the membrane receptor complex of the Wnt/β-catenin pathway and with the vacuolar-type H-ATPase (15),(16).

Activation of the RAS in obesity hypertension stems from various factors. Apart from the fact that SNS activation and the compression of the kidneys activate the RAS, components of the RAS are themselves activated from what seems to be an impaired negative feedback loop (4, 5). Additionally, elevated tissue RAS in tissues such as adipose tissue, not only perpetuates hypertension pathophysiology but also contributes to obesity. Ang II is known to exert a trophic effect on adipocyte differentiation and adipose tissue growth which is mediated by Angiotensin type 1 receptor (AT1R) and ERK1/2 activation (17, 18).

PRR and sPRR have also been shown to play a role in the developing obesity and obesity related hypertension. Knockdown of adipose specific PRR and whole-body loss of sPRR reduced body weight in mice (19–22). Moreover, obesity is known to increase adipose tissue PRR levels (19, 23) and elevate circulating sPRR (24) whereas weight loss after bariatric surgery decreased circulating sPRR in both men and women (24). However, a recent study in mice showed that elevated plasma sPRR could improve severe obesity and its comorbidities, such as hyperlipidemia (25).

Previous studies have shown that there is a role of sPRR during obesity. Specifically, infusion of mouse recombinant sPRR increased systolic blood pressure (SBP) and impaired the baroreflex sensitivity likely through the sympatho-excitatory effects of leptin on blood pressure in high fat (HF)-fed male C57BL/6J mice (26). Conversely, loss of sPRR decreased blood pressure and attenuated Ang II-induced hypertension (22). This data indicates that elevated levels of circulating mouse sPRR display pro-hypertensive effects and mediate the pressor effects of Ang II.

Clinical studies have previously shown that elevated circulating sPRR levels is associated with renal dysfunction (27, 28) heart failure (27) essential hypertension (29) and with obstructive sleep apnea in obese patient (30). In patients with type 2 diabetes, plasma sPRR correlated with activation of systemic RAS (elevated plasma renin activity) in women whiles urine sPRR correlated with intrarenal RAS (elevated urinary renin activity) activation in men (31).

In the kidney, expression of PRR has been shown to increase in diabetes and high glucose feeding leading to kidney injury (32, 33); yet intrarenal PRR and sPRR effects on blood pressure during obesity are poorly understood. Similarly, the role of renal-derived human sPRR (HsPRR) in kidney function and blood pressure regulation during obesity has not been investigated. Therefore, this study sought to assess the contribution of human sPRR derived from the kidney on blood pressure and renal hemodynamics. For that, we used a mouse model that expresses human sPRR in the collecting duct (CD-HsPRR) to determine the impact on blood pressure, GFR, and water and electrolyte homeostasis, while correlating blood pressure with renal cortex RAS and ion transporters

## Methods

The original data that support the findings of this study are available from the corresponding author upon reasonable request.

### Transgenic mouse model

All animal studies described below complied with the University of Kentucky Institutional Animal Care and Use Committee (Protocol No. 2022-4059) and followed the “Guide for the Care and Use of Laboratory Animals” issued by the National Research Council. Mice were on a 14:10 hours, light: dark cycle, starting at 7:00 am for the light cycle and at 9:00 pm for the dark cycle. Mice had *ad libitum* access to water and diet.

Human sPRR-Myc-tag transgenic mice were developed by cloning myc epitope-tagged sPRR into the CAG-GFP vector (34). The fragment was sequenced to confirm accuracy of the construct and transfected into HEK293 cells stably expressing Cre recombinase and screened with a myc monoclonal antibody (clone Vli01, Maine Health Institute for Research, Scarborough, ME) to verify the expression of the fragment in the media. Upon validation, the construct was then injected into oocytes of C57BL/6J mice to generate male and female Human sPRR-Myc-tagged transgenic mice.

CD-HsPRR male and female mice were generated by breeding heterozygote female human sPRR-Myc-tag transgenic mice with male mice expressing Cre recombinase under the control of the HoxB7 promoter (STOCK Tg (Hoxb7-cre)13Amc/J, #004692, The Jackson Laboratory, Figure 1A). HoxB7/Cre transgenic mice have been previously used to specifically target renal collecting duct cells (35).

**Figure 1.**
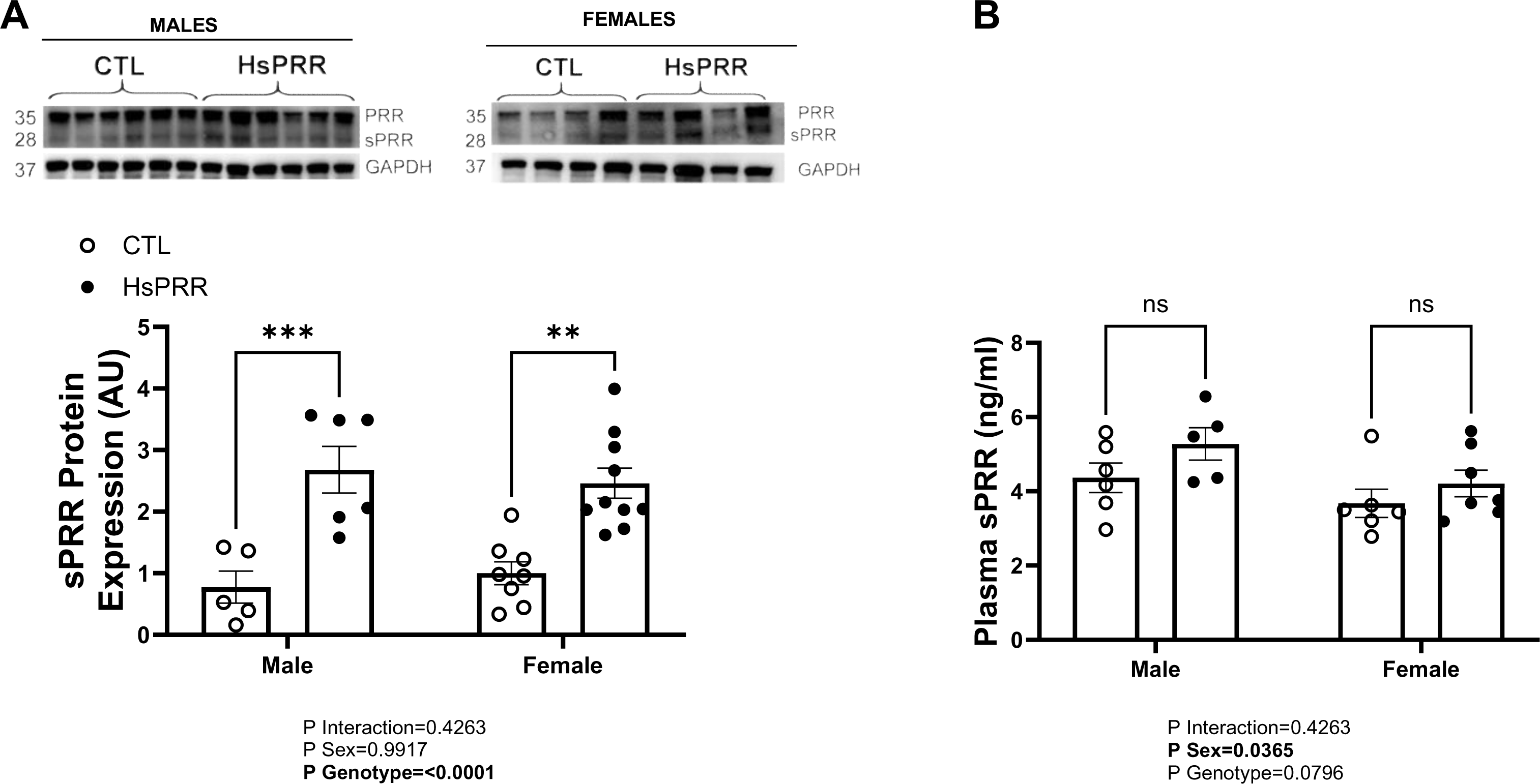
Tissue and plasma sPRR in CD-HsPRR during obesity. Protein expression of sPRR in kidney male and female collecting duct derived HsPRR mice (Figure 1A). Plasma sPRR in male and female mice (Figure 1B). Data are expressed as mean ± SEM of 6-8 mice/group. A 2-way ANOVA was performed to detect differences *P<0.05 compared with CTL.

### Experimental Design

CD-HsPRR and control littermates (CTL) male and female mice (CTL= 9 males and 9 males; K-HsPRR= 8 males and 7 females). Mice were placed on a regular chow at weaning and switched to a high fat diet (HFD: 60 % kcal from fat; D12492, Research Diets, Inc., USA) for 25 weeks. Mice had *ad libitum* access to water and diet. Body weight was recorded weekly. At 10 weeks of HFD, the glomerular filtration rate was assessed and at 12 weeks, mice were placed individual metabolic cages (Techniplast Solo Mouse Metabolic Cages; Techniplast USA, Exton, PA) with free access to food and water for urine collection tube was coated with mineral oil to prevent evaporation (20).

At 20 weeks, mice were housed individually and implanted in the left carotid artery with a catheter connected to a telemetry transmitter for acute blood pressure measurements as described in Chapter 2, General Methods. At the end of the study, tissues and plasma were collected for biochemical analysis.

### Glomerular filtration rate measurement

The glomerular filtration rate was measured at week 10 of the study as described previously (36). Mice were placed under light anesthesia (isoflurane) and the flank area was depilated to apply the transcutaneous receiver on top (NIC-kidney device). The receiver was secured around the body with 3M surgical tape. After the animal awakened from anesthesia, a 5-min baseline trace was recorded. Then, they were injected with 30 μl of fluorescein isothiocyanate-sinistrin (FITC-sinistrin) retro-orbitally (5 mg/100 g BW in 0.9% saline, Fresenius Kabi, Linz, Austria) under light isoflurane using microneedles. After 90 minutes of measurement, the device was removed. The probe was read to determine t_1/2_ in minutes. Renal function was evaluated by the elimination kinetics of fluorescence decay (three-compartmental model) using the following formula=14616.8/(t_1/2_) = GFR (μl/min/100 g BW).

### Western Blot Analysis

Protein from frozen kidney was extracted in ice-cold RIPA buffer (Thermo Scientific, Waltham, MA) with a Geno/Grinder 2010 (SPEX Sample Prep, Metuchen, NJ). Protein obtained was run on SDS-PAGE on precast polyacrylamide gel (Mini-PROTEAN TGX, 4%-20%; Bio-Rad Laboratories, Hercules, CA) and transferred to a polyvinylidene difluoride membrane via a semi-wet transfer equipment (Trans-Blot Turbo Transfer System; Bio-Rad Laboratories, Hercules, CA) as recommended by the manufacturer. Membranes were blocked in 5% nonfat dried milk in Tris-buffered saline with 0.1% Tween 20, membranes were incubated with C-myc antibody (MaineHealth Institute for Research, clone Vli01), Anti-ATP6AP2 (Millipore Sigma, HPA003156), Aquaporin 2 (Cell Signaling Technology Inc, 3487) and GAPDH (Sigma Aldrich, G9545) in 5% nonfat dried milk in Tris-buffered saline with 0.1% Tween 20. After incubation with handle region decoy peptide-conjugated antirabbit secondary antibody (Jackson ImmunoResearch Laboratories, AB2313567), proteins were imaged using ChemiDoc Imaging Sysytem (Bio-Rad Laboratories, Hercules, CA). The levels of proteins were quantified using Image Lab software (Bio-Rad Laboratories, Hercules, CA) and normalized to GAPDH.

### Tissue RNA Extraction and Quantitative RT-PCR

RNA from the kidney was extracted with the RNeasy Fibrous Tissue Mini Kit (Qiagen, Madison, WI). Kidney cortex samples were first lysed in Buffer RLT and then diluted before being treated with proteinase K. Debris was pelleted by centrifugation at room temperature for 3 minutes, and the supernatant was removed. The supernatant was mixed with ethanol and then centrifuged through a RNeasy Mini spin column, where RNA binds to the silica membrane. Traces of DNA that may co-purify with the RNA were removed by DNase treatment on the silica membrane. DNase and any contaminants are efficiently washed away, and high-quality total RNA was eluted in RNase-free water. RNA concentration was measured by a NanoDrop (Thermo-Fisher Scientific, Madison, WI), and the cDNA was synthesized using Qscript cDNA SuperMix (Quanta Biosciences, Gaithersburg, MD). mRNA expression of genes (Supplemental Table 3) was evaluated by quantitative RT-PCR using PerfeCTa SYBR Green FastMix (Quanta Biosciences) and normalized to GAPDH. Thermocycling protocol consisted of 5 min at 95C, 40 cycles of 15 s at 95C, and 30 s at 60C, finishing with a melting curve ranging from 60 to 95C to allow distinction of specific products.

### Statistical Analysis

Data are represented as means±SEM. Statistical analysis was performed using Graph Prism. Statistical differences between groups were assessed by a two-way, ANOVAS followed by Tukey post hoc analysis for multiple comparisons. Grubbs test (GraphPad QuickCalcs) was used to determine statistical outliers. Values of P<0.05 were considered statistically significant.

## Results

### CD-HsPRR increases intrarenal but not plasma sPRR

Body and kidney weights were similar between groups and sexes (Supplementary Table 1). CD-HsPRR display increased intrarenal sPRR but did not change circulating sPRR levels when fed an HFD for 25 weeks (Figure 1A-B).

### High fat diet increases blood pressure similarly in mice

Blood pressure was similar between CTL and CD-HsPRR mice in males and females at both daytime and nighttime (Figure 2A-F). Heart rate was also similar between groups in both males and females (Figure 2G-H). Hence, no changes were observed at 24 hours for blood pressure and heart (Supplementary Table 2).

**Figure 2.**
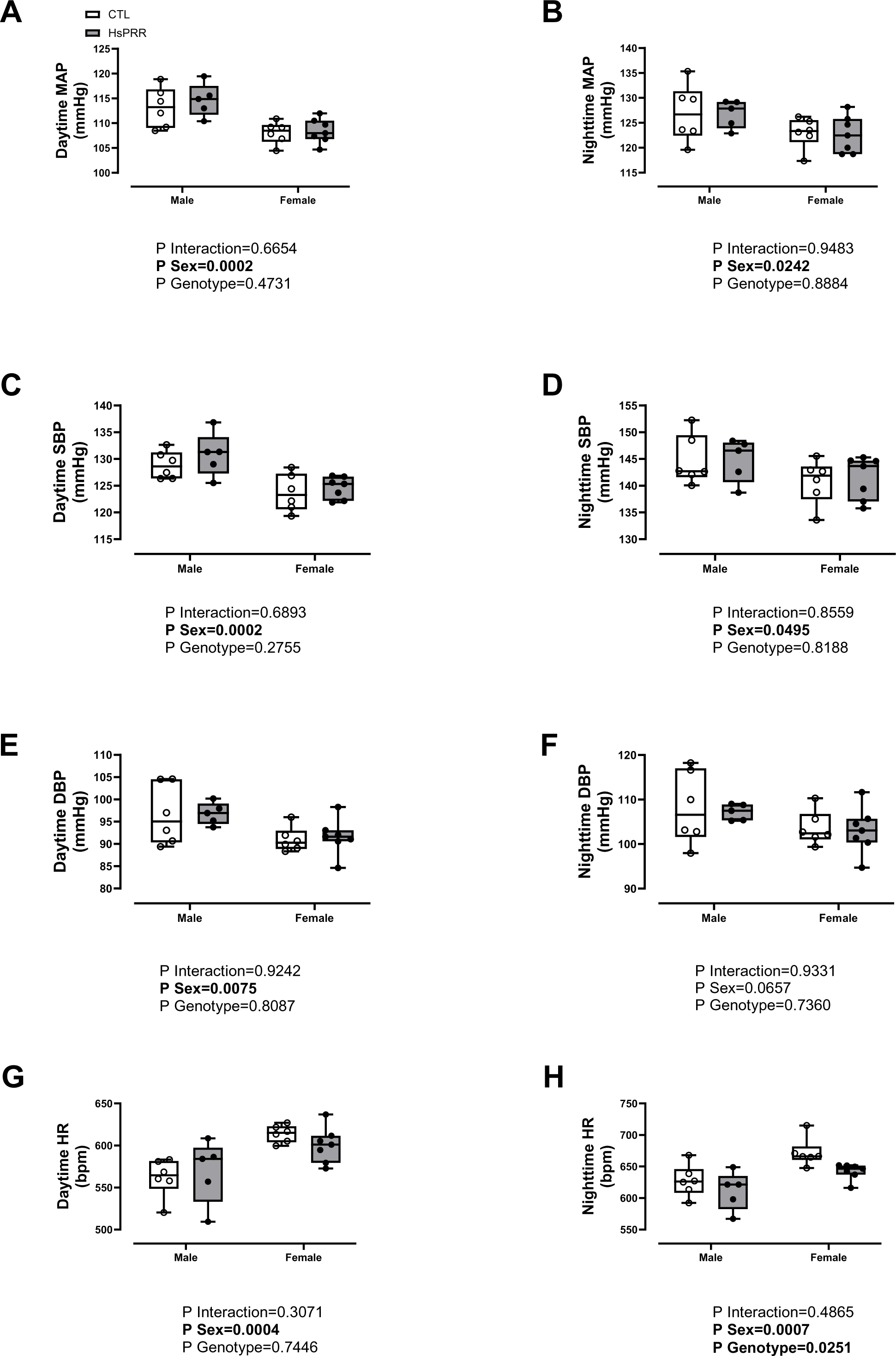
Blood pressure outcomes in male and female mice. Mean arterial blood pressure at (A) daytime and (B) nighttime; Systolic blood pressure at (C) daytime and (D) nighttime; Diastolic blood pressure at (E) daytime and (F) nighttime; Heart rate at (F) daytime and (G) nighttime. Data are expressed as mean ± SEM of 6-9 mice/group. A two-way ANOVA was performed to detect differences. * P<0.05 compared with CTL.

### Obesity leads to losartan resistance in CD-HsPRR male mice

Next, we investigated the influence of CD-HsPRR on pressor response to acute Ang II, which was similar in both male and female mice (Figure 3A). Also, we blocked various receptors mediating the AngII (losartan), sympathetic (chlorisondamine) and parasympathetic (atropine) signaling. Losartan-induced antihypertensive effect was present was impaired only in male CD-HsPRR mice (Figure 3B). Moreover, a ganglion blocker and a parasympathetic antagonist showed similar changes in blood pressure in CTL and K-HsPRR mice (Figure 3C-D).

**Figure 3.**
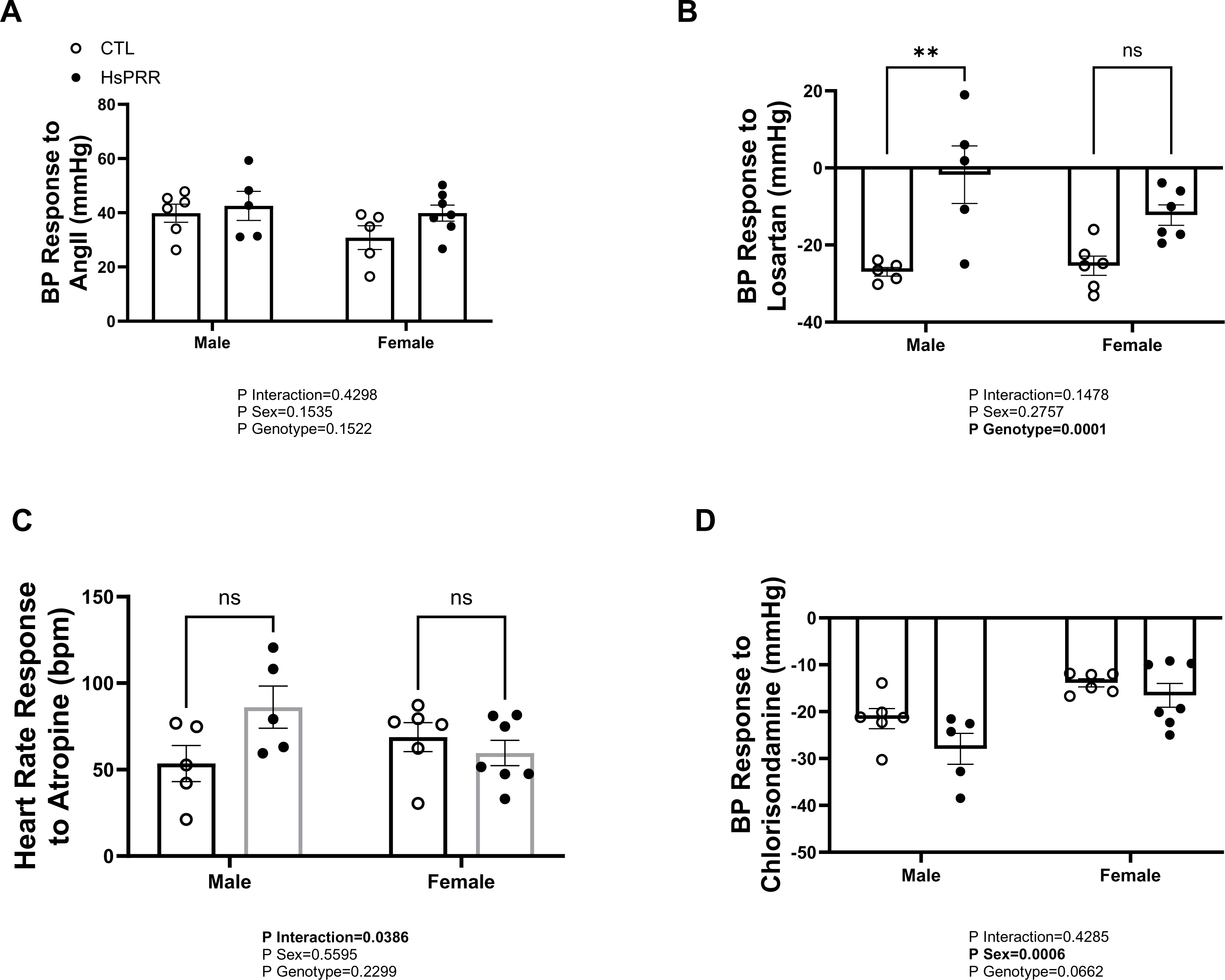
CD-HsPRR blunts BP response to losartan in male mice. (A) Acute blood pressure response (5 mins) to Ang II injection in K-HsPRR mice. (B) Peak blood pressure response (3 h) to losartan injection. (C) Heart rate response (1 h) to atropine injection (D) Blood pressure response (1 h) to Chlorisondamine in K-HsPRR mice. Data are expressed as mean ± SEM of 5-7 mice/group. A two-way ANOVA was performed to detect differences in. * P<0.05 compared with CTL.

### CD-HsPRR does not affect GFR but upregulates ENaC expression

The CD-HsPRR expression did not change glomerular filtration rate (GFR), water intake or urine flow in both male and female mice (Fig. 4A-C). However, the water transporter, AQP2 and sodium channel, β- and γ-ENaC were upregulated in the male CD-HsPRR group (Fig. 5A, C, D) while α-ENaC protein expression was decreased in females CD-HsPRR (Fig. 5B).

**Figure 4.**
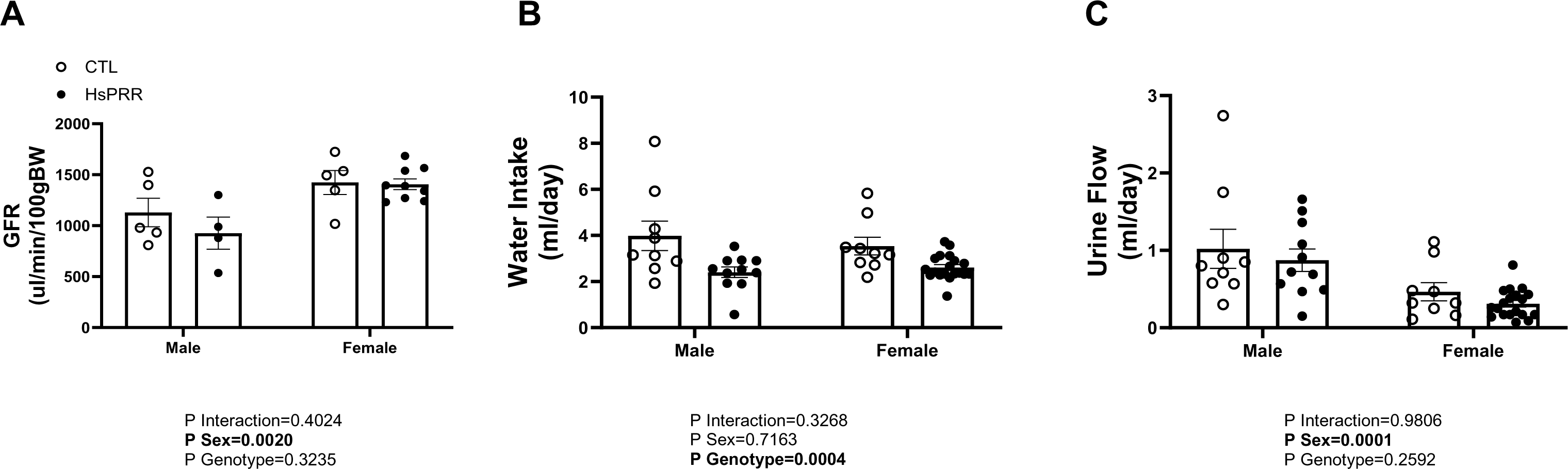
CD-HsPRR does not affect GFR and fluid homeostasis during obesity. (A) Glomerular filtration rate (GFR). Data are expressed as mean ± SEM of 4-9 mice/group (B) Water intake; (C) Urine flow rate. Data are expressed as mean ± SEM of 9-17 mice/group; A two-way ANOVA was performed to detect differences. * P<0.05 compared with CTL.

**Figure 5.**
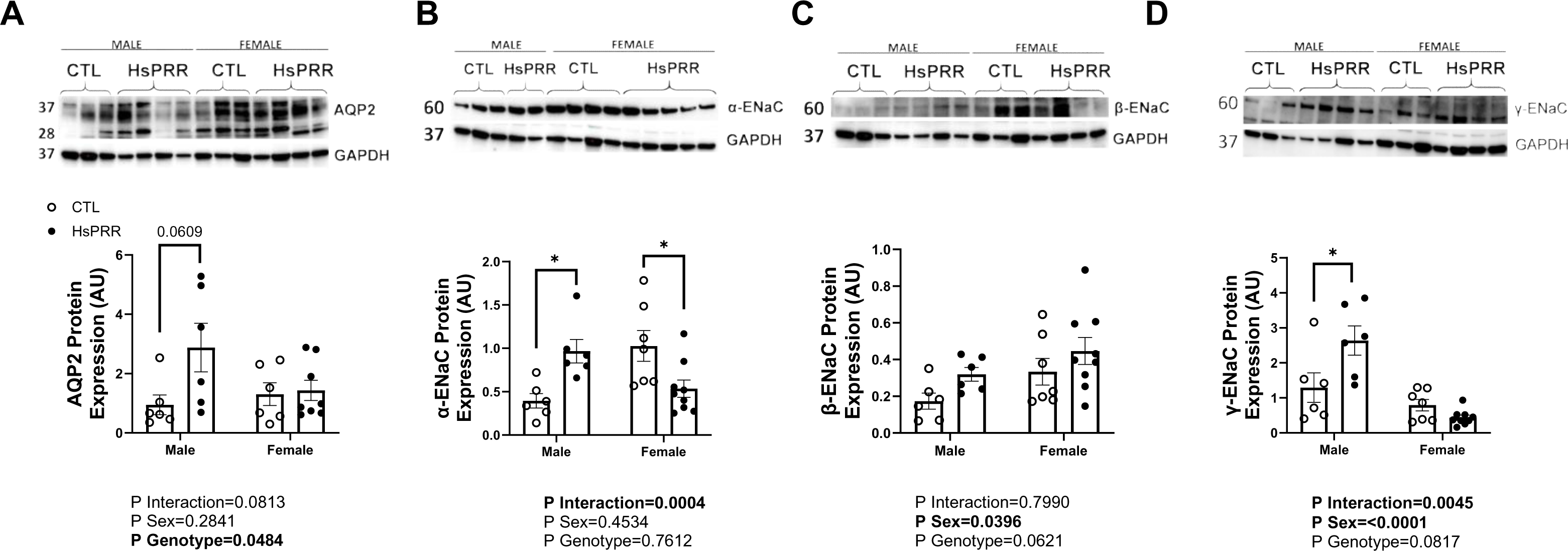
Renal-derived human sPRR modulates α- and β-ENaC expression during obesity. (A) Aquaporin2 protein expression; (B) Alpha-ENaC protein expression; (C) Beta-ENaC protein expression and (D) Gamma-ENaC protein expression in the kidney. Data are expressed as mean ± SEM of 6-8 mice/group. A two-way ANOVA was performed to detect differences. * P<0.05 compared with CTL.

### CD-HsPRR upregulates intrarenal renin and AT1R in males during obesity

To determine whether CD-HsPRR modulates intrarenal RAS expression, we assessed the mRNA expression of AGT, renin, AT1R, and ACE in the kidney cortex. When comparing CLT and CD-HsPRR in male mice, AGT and ACE mRNA abundance were similar between groups, while renin and AT1R expression was upregulated (Figure 6A). All four RAS components were similar between CTL and female CD-HsPRR mice (Figure 6B). COX2 expression was not influenced by the expression of HsPRR in both sexes (Figure 6C).

**Figure 6.**
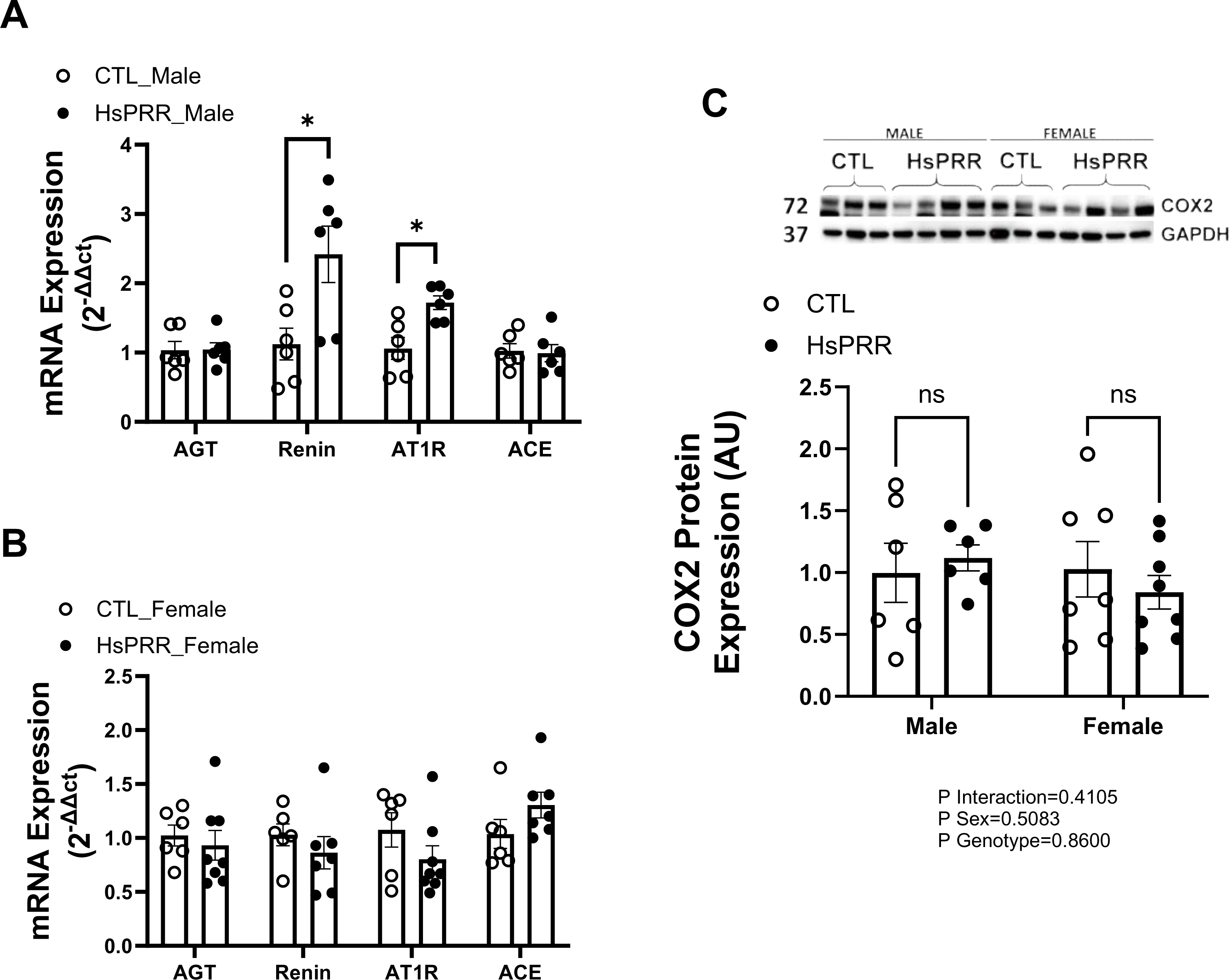
CD-HsPRR modulates expression of renin and AT1R in the kidney. Renal RAS gene expression in kidney of (A) Male and (B) Female CD-HsPRR mice; Data are expressed as mean ± SEM of 6-8 mice/group. A t-test was performed to detect differences.

### Obesity increases plasma leptin in female CD-HsPRR mice

CD-HsPRR expression did not affect the body weight gain between groups; however, increased gonadal white adipose tissue (gWAT) in female CD-HsPRR mice only (Figure 7A). Along these lines, plasma leptin was increased near 3-fold in female CD-HsPRR mice (Figure 7B).

**Figure 7.**
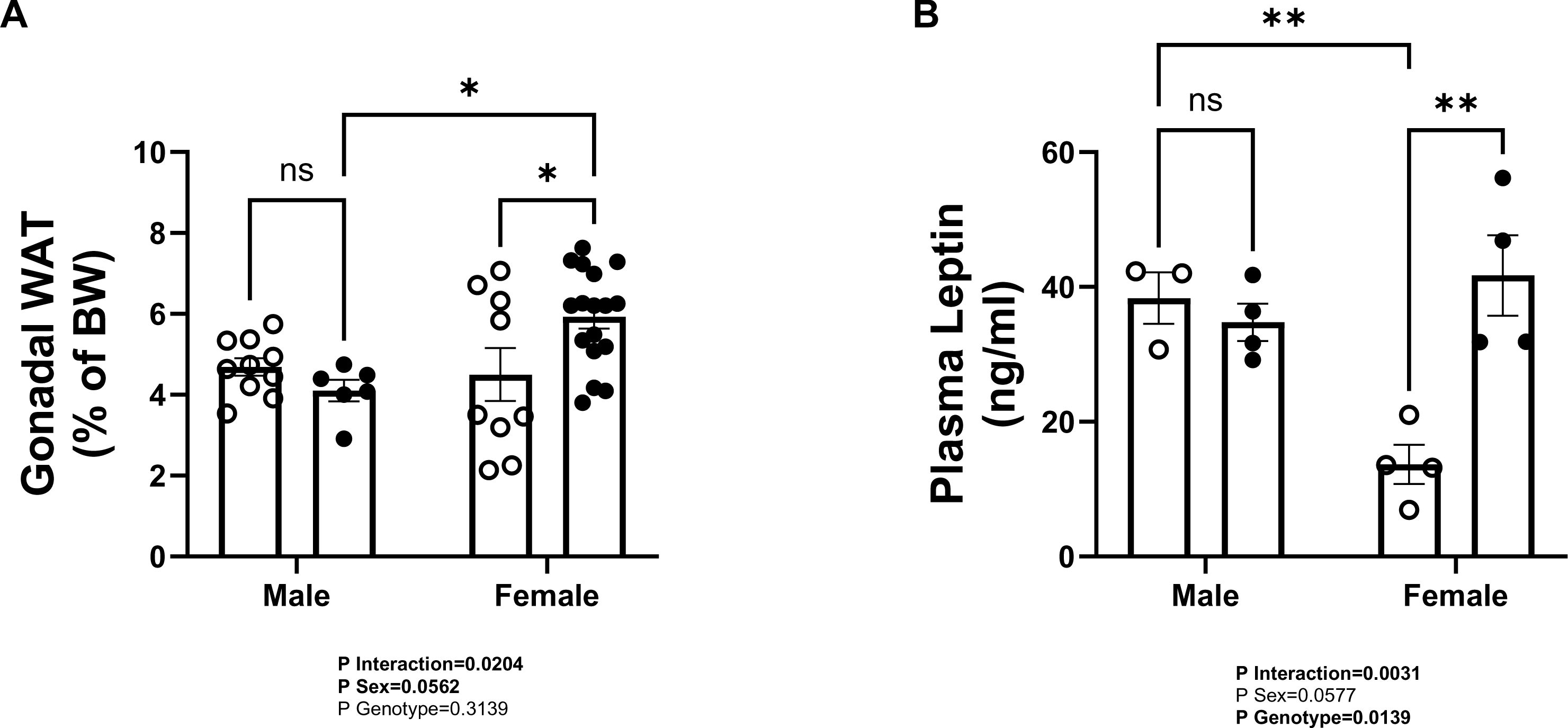
CD-HsPRR increases plasma leptin in CD-HsPRR mice. Gonadal white adipose tissue. Data are expressed as mean ± SEM of 6-17 mice/group (B) Plasma leptin. Data are expressed as mean ± SEM of 3-5 mice/group. A two-way ANOVA was performed to detect differences. * P<0.05 compared with CTL

## Discussion

The study aimed to determine whether obesity driven by a high fat diet would increase and/or exacerbate blood pressure and impair renal function outcomes observed at physiological conditions. Using a novel mouse model that expresses human sPRR in the collecting duct (CD) of the kidney, the current study reports the following findings in CD-HsPRR mice fed a HFD compared to CTL mice: (1) unchanged blood pressure (2) pressor response to a single dose of losartan impaired only in males; (3) unchanged GFR (4) increased α-and ENaC expression in males but decreased in female mice; increased gonadal WAT and plasma leptin in female mice; (6) increased renin expression in male mice without changing COX2 expression in renal cortex. Furthermore, intrarenal sPRR expression was increased in CD-HsPRR mice while plasma sPRR remained similar to controls, while water intake and urine flow and AQP2 expression were not influenced by HsPRR expression in male and female mice. Taken together, our study shows that the expression of HsPRR in CD cells does not increase blood pressure further in response to an obesogenic diet despite intrarenal changes in gene and protein expression that are associated with the water and electrolyte homeostasis, indicating that there are antihypertensive mechanisms associated to HFD that could buffer the effect on sodium reabsorption and vascular resistance.

In humans, obesity is associated with increased plasma sPRR levels, whereas bariatric surgery can normalize these concentrations (24). This data suggests that fat mass is directly proportional to the circulating levels of sPRR. Our study shows that although intrarenal sPRR expression was increased in male and female mice, plasma sPRR was unchanged compared with CTL mice. This result reinforces the hypothesis that the expression of HsPRR in the kidney does not influence the circulating sPRR levels. We studied how renal expression of HsPRR influenced body composition in response to HFD. We found that gonadal WAT and plasma leptin were increased in female CD-HsPRR mice, along with proportional increases in adipose tissue mass, a well-known factor contributing obesity-associated hypertension (37). In the kidney, the role of leptin in renal hemodynamics is still unclear; however, accumulating evidence shows that leptin is involved in sodium balance. For instance, in mice treated with leptin for a short term (7 days), leptin induced an increase in filtration faction and natriuresis with no change in GFR. Further, long-term leptin treatment (28 days) led to a decrease in renal plasma flow and filtration fraction without any change in GFR or tubular function (38). Along these lines, rats treated with human leptin acutely showed increased natriuretic activity without any change in the arterial pressure (39). Moreover, elevated leptin, either systemically or locally in the kidney reduces Na+-K+-ATPase activity in the renal medulla (40–42). Finally leptin receptors are predominant in the collecting duct (43), suggesting that most of the leptin-induced changes are most likely mediated by tubular mechanisms. Together, these studies show that leptin can modulate sodium handling by increasing sodium excretion. Accordingly, decreased α-ENaC expression in female CD-HsPRR fed an HFD could increase natriuretic activity, lowering blood pressure and normalizing GFR. This idea is further supported by the impaired response to obesity-induced hypertension only observed in female HsPRR mice (see general discussion).

Another possibility that prevents high blood pressure in females fed a high fat diet similar to what we have observed with regular chow may be because obesity has been shown to increase plasma estrogen levels (44). Hence, a likely increase in estrogen due to the exacerbated visceral adiposity in female CD-HsPRR mice could have prevented the activation of the intrarenal RAS and increased blood pressure. Indeed, female CD-HsPRR mice fed a HFD did not have any change in RAS expression in the kidney cortex as the ones fed regular chow (Chapter 3).

Paradoxically, clinical studies show that elevated leptin leads to renal dysfunction particularly in women (45). Moreover, other studies examining the impact of leptin on kidney function have shown an anti-natriuretic action and renal damage (46–48). Leptin is also known to increase blood pressure in females via an aldosterone-dependent mechanism (49–51). These discrepancies reported in the literature show that further studies are warranted to understand the mechanism and counteractive pathways in leptin-mediated sodium handling, renal injury, and kidney-dependent hypertension in the presence of elevated local levels of HsPRR/sPRR.

In obese males, although blood pressure was similar between groups, CD-HsPRR elicited resistance to lower blood pressure in response to a dose of losartan. Additionally, obese male CD-HsPRR mice have an increase in α-and ENaC protein expression along with renin and AT1R mRNA expression. The role of sPRR in obesity-related hypertension in male mice has been well demonstrated (52–54). Male mice fed a high fat diet for 32 weeks showed elevated plasma sPRR was associated with activation of systemic RAS, vascular damage, and non-dipping hypertension (54). Similarly, obese male mice fed 32 weeks of HFD and infused with sPRR had hypertension due to endothelial dysfunction mediated by the activation of AT1R (52). sPRR infusion has been shown to increase blood pressure in male mice via a leptin-dependent pathway (53). In the current study, HFD feeding for 25 weeks in male mice expressing human sPRR in the collecting duct did not elevate plasma sPRR, and plasma leptin nor elicit an increase in blood pressure. Possible reasons for the discrepancies could be the length of feeding (25 weeks vs 32 weeks) and the unchanged levels of circulating sPRR. Obese male CD-HsPRR mice, however, showed impaired losartan response and increased intrarenal renin and AT1R mRNA expression, indicating that HsPRR in the kidney can impact the RAS-dependent mechanisms implicated in lowering blood pressure. Moreover, CD-HsPRR increased expression of sodium channel α and - NaC, which could be secondary to the intrarenal activation of RAS. However, there were no changes in water intake, urine flow, and GFR, suggesting that a possible negative feedback loop such as increase in potassium reabsorption, could be present to counteract the effect of a likely increase in luminal and plasma sodium. Future studies are warranted to elucidate the impact of HsPRR in sodium handling during obesity in males and possible feedback loops. Taken together, our study shows that although CD-HsPRR in obesity does not elevate blood pressure in both males and females, it may contribute to BP control mechanisms in males via the AT1R. Thus, renal sPRR could be a potential therapeutic target for resistant hypertension in obese males.

## Acknowledgments

We thank the Mouse Transgenic Core (supported by COBRE 1P20GM121301, L. Liaw PI) at the MaineHealth Institute for Research for generation of the Cre-inducible human sPRR-myc-tag transgenic mouse strain.

We thank Dr. Wen Su from Dr. Ming Gong Laboratory for her assistance with radiotelemetry surgery.

## Disclaimers and Disclosures

None

## Supplemental Materials

T1-T2

## REFERENCES

1. Organization WH. Obesity and overweight https://www.who.int/news-room/fact-sheets/detail/obesity-and-overweight2021 [Available from: https://www.who.int/news-room/fact-sheets/detail/obesity-and-overweight.

2. Steirman B. National Health and Nutrition Examination Survey 2017 - March 2020 prepandemic data files : development of files and prevalence estimates for selected health outcomes. Hyattsville, MD: U.S. Department of Health and Human Services, Centers for Disease Control and Prevention, National Center for Health Statistics; 2021.

3. Alpert MA, Omran J, Mehra A, Ardhanari S. Impact of obesity and weight loss on cardiac performance and morphology in adults. Prog Cardiovasc Dis. 2014;56(4):391–400.

4. Hall ME, Cohen JB, Ard JD, Egan BM, Hall JE, Lavie CJ, et al. Weight-Loss Strategies for Prevention and Treatment of Hypertension: A Scientific Statement From the American Heart Association. Hypertension. 2021;78(5).

5. Shariq OA, Mckenzie TJ. Obesity-related hypertension: a review of pathophysiology, management, and the role of metabolic surgery. Gland Surgery. 2020;9(1):80–93.

6. Jiang S-Z, Lu W, Zong X-F, Ruan H-Y, Liu Y. Obesity and hypertension. Experimental and Therapeutic Medicine. 2016;12(4):2395–9.

7. Putnam K, Shoemaker R, Yiannikouris F, Cassis LA. The renin-angiotensin system: a target of and contributor to dyslipidemias, altered glucose homeostasis, and hypertension of the metabolic syndrome. Am J Physiol Heart Circ Physiol. 2012;302(6):H1219–30.

8. Szczepanska-Sadowska E, Czarzasta K, Cudnoch-Jedrzejewska A. Dysregulation of the Renin-Angiotensin System and the Vasopressinergic System Interactions in Cardiovascular Disorders. Curr Hypertens Rep. 2018;20(3):19.

9. Ichihara A, Yatabe MS. The (pro)renin receptor in health and disease. Nat Rev Nephrol. 2019;15(11):693–712.

10. Nguyen G, Muller DN. The biology of the (pro)renin receptor. J Am Soc Nephrol. 2010;21(1):18–23.

11. Nabi AH, Kageshima A, Uddin MN, Nakagawa T, Park EY, Suzuki F. Binding properties of rat prorenin and renin to the recombinant rat renin/prorenin receptor prepared by a baculovirus expression system. Int J Mol Med. 2006;18(3):483–8.

12. Nguyen G, Burckle C, Sraer JD. [Proteases and antiproteases in the progression of chronic renal insufficiency lesions. The role of the tissue renin-angiotensin system and the renin receptor]. J Soc Biol. 2002;196(4):281–4.

13. Nguyen G, Delarue F, Burckle C, Bouzhir L, Giller T, Sraer JD. Pivotal role of the renin/prorenin receptor in angiotensin II production and cellular responses to renin. J Clin Invest. 2002;109(11):1417–27.

14. Sakoda M, Ichihara A, Kaneshiro Y, Takemitsu T, Nakazato Y, Nabi AH, et al. (Pro)renin receptor-mediated activation of mitogen-activated protein kinases in human vascular smooth muscle cells. Hypertens Res. 2007;30(11):1139–46.

15. Ludwig J, Kerscher S, Brandt U, Pfeiffer K, Getlawi F, Apps DK, et al. Identification and characterization of a novel 9.2-kDa membrane sector-associated protein of vacuolar proton-ATPase from chromaffin granules. J Biol Chem. 1998;273(18):10939–47.

16. Cruciat CM, Ohkawara B, Acebron SP, Karaulanov E, Reinhard C, Ingelfinger D, et al. Requirement of prorenin receptor and vacuolar H+-ATPase-mediated acidification for Wnt signaling. Science. 2010;327(5964):459-63.

17. Darimont C, Vassaux G, Ailhaud G, Negrel R. Differentiation of preadipose cells: paracrine role of prostacyclin upon stimulation of adipose cells by angiotensin-II. Endocrinology. 1994;135(5):2030–6.

18. Saint-Marc P, Kozak LP, Ailhaud G, Darimont C, Negrel R. Angiotensin II as a trophic factor of white adipose tissue: stimulation of adipose cell formation. Endocrinology. 2001;142(1):487–92.

19. Shamansurova Z, Tan P, Ahmed B, Pepin E, Seda O, Lavoie JL. Adipose tissue (P)RR regulates insulin sensitivity, fat mass and body weight. Mol Metab. 2016;5(10):959–69.

20. Gatineau E, Cohn D, Poglitsch M, Loria A, Gong M, Yiannikouris F. Losartan prevents the elevation of blood pressure in adipose-PRR deficient female mice while elevated circulating sPRR activates the renin-angiotensin system. American Journal of Physiology. 2019;316(3):H506.

21. Wu C-H, Mohammadmoradi S, Thompson J, Su W, Gong M, Nguyen G, et al. Adipocyte (Pro)Renin-Receptor Deficiency Induces Lipodystrophy, Liver Steatosis and Increases Blood Pressure in Male Mice. Hypertension. 2016;68(1):213–9.

22. Ramkumar N, Stuart D, Peterson CS, Hu C, Wheatley W, Cho J-M, et al. Loss of Soluble (Pro)renin Receptor Attenuates Angiotensin-II Induced Hypertension and Renal Injury. Circulation research. 2021.

23. Achard V, Boullu-Ciocca S, Desbriere R, Nguyen G, Grino M. Renin receptor expression in human adipose tissue. Am J Physiol Regul Integr Comp Physiol. 2007;292(1):R274–82.

24. Nishijima T, Ohba K, Baba S, Kizawa T, Hosokawa K, Endo F, et al. Decrease of Plasma Soluble (Pro)renin Receptor by Bariatric Surgery in Patients with Obstructive Sleep Apnea and Morbid Obesity. Metabolic syndrome and related disorders. 2018;16(4):174–82.

25. Wang F, Luo R, Zou C-J, Xie S, Peng K, Zhao L, et al. Soluble (pro)renin receptor treats metabolic syndrome in mice with diet-induced obesity via interaction with PPARγ. JCI insight. 2020;5(7).

26. Gatineau E, Gong MC, Yiannikouris F. Soluble Prorenin Receptor Increases Blood Pressure in High Fat-Fed Male Mice. Hypertension. 2019;74(4):1014–20.

27. Fukushima A, Kinugawa S, Homma T, Masaki Y, Furihata T, Abe T, et al. Increased plasma soluble (pro)renin receptor levels are correlated with renal dysfunction in patients with heart failure. Int J Cardiol. 2013;168(4):4313–4.

28. Hamada K, Taniguchi Y, Shimamura Y, Inoue K, Ogata K, Ishihara M, et al. Serum level of soluble (pro)renin receptor is modulated in chronic kidney disease. Clin Exp Nephrol. 2013;17(6):848–56.

29. Morimoto S, Ando T, Niiyama M, Seki Y, Yoshida N, Watanabe D, et al. Serum soluble (pro)renin receptor levels in patients with essential hypertension. Hypertens Res. 2014;37(7):642–8.

30. Nishijima T, Ohba K, Baba S, Kizawa T, Hosokawa K, Endo F, et al. Decrease of Plasma Soluble (Pro)renin Receptor by Bariatric Surgery in Patients with Obstructive Sleep Apnea and Morbid Obesity. Metab Syndr Relat Disord. 2018;16(4):174–82.

31. Visniauskas B, Arita DY, Rosales CB, Feroz MA, Luffman C, Accavitti MJ, et al. Sex differences in soluble prorenin receptor in patients with type 2 diabetes. Biology of Sex Differences. 2021;12(1).

32. Helmy MS, Jiqian H. Renal (pro)renin receptor upregulation in diabetic rats through enhanced angiotensin AT1 receptor and NADPH oxidase activity. Experimental Physiology. 2008;93(5):709–14.

33. Li C, Siragy HM. High Glucose Induces Podocyte Injury via Enhanced (Pro)renin Receptor-Wnt-β-Catenin-Snail Signaling Pathway. PloS one. 2014;9(2):e89233-e.

34. Duarte CW, Stohn JP, Wang Q, Emery IF, Prueser A, Lindner V. Elevated Plasma Levels of the Pituitary Hormone Cthrc1 in Individuals with Red Hair but Not in Patients with Solid Tumors. PLoS ONE. 2014;9(6):e100449.

35. Prieto M, Reverte V, Mamenko M, Kuczeriszka M, Veiras L, Rosales C, et al. Collecting duct prorenin receptor knockout reduces renal function, increases sodium excretion, and mitigates renal responses in ANG II-induced hypertensive mice. American Journal of Physiology. 2017;313(6):F1243.

36. Ellery SJ, Cai X, Walker DD, Dickinson H, Kett MM. Transcutaneous measurement of glomerular filtration rate in small rodents: Through the skin for the win?: Transcutaneous measure of GFR in rodents. Nephrology (Carlton, Vic). 2015;20(3):117–23.

37. Lundgren M, Svensson M, Lindmark S, Renstrom F, Ruge T, Eriksson JW. Fat cell enlargement is an independent marker of insulin resistance and ’hyperleptinaemia’. Diabetologia. 2007;50(3):625–33.

38. Thieme K, Oliveira-Souza M. Renal Hemodynamic and Morphological Changes after 7 and 28 Days of Leptin Treatment: The Participation of Angiotensin II via the AT1 Receptor. PLOS ONE. 2015;10(3):e0122265.

39. Jackson EK, Li P. Human leptin has natriuretic activity in the rat. American Journal of Physiology - Renal Physiology. 1997;272(3):333–8.

40. Bettowski J, G Wj, Górny D, Marciniak A. Human leptin administered intraperitoneally stimulates natriuresis and decreases renal medullary Na+, K+-ATPase activity in the rat -- impaired effect in dietary-induced obesity. Med Sci Monit. 2002;8(6):Br221–9.

41. Bettowski J, Wojcicka G. Spectrophotometric method for the determination of renal ouabain-sensitive H+,K+-ATPase activity. Acta Biochim Pol. 2002;49(2):515–27.

42. Bettowski J, Marciniak A, Wójcicka G. Leptin decreases renal medullary Na(+), K(+)-ATPase activity through phosphatidylinositol 3-kinase dependent mechanism. J Physiol Pharmacol. 2004;55(2):391–407.

43. Martrnez-Anso E, Lostao MP, Martinez JA. Immunohistochemical localization of leptin in rat kidney. Kidney Int. 1999;55(3):1129–30.

44. Cleary MP, Grossmann ME. Minireview: Obesity and breast cancer: the estrogen connection. Endocrinology. 2009;150(6):2537–42.

45. Pedone C, Roshanravan B, Scarlata S, Patel KV, Ferrucci L, Incalzi RA. Longitudinal association between serum leptin concentration and glomerular filtration rate in humans. PloS one. 2015;10(2):e0117828-e.

46. Bettowski J, Jamroz-Wiśniewska A, Borkowska E, Wójcicka G. Up-regulation of renal Na+, K+-ATPase: the possible novel mechanism of leptin-induced hypertension. Pol J Pharmacol. 2004;56(2):213–22.

47. Beltowski J, Wójcicka G, Marciniak A, Jamroz A. Oxidative stress, nitric oxide production, and renal sodium handling in leptin-induced hypertension. Life Sci. 2004;74(24):2987–3000.

48. Wolf G, Chen S, Han DC, Ziyadeh FN. Leptin and renal disease. American Journal of Kidney Diseases. 2002;39(1):1–11.

49. Huby A-C, Otvos L, Belin de Chantemèle EJ. Leptin Induces Hypertension and Endothelial Dysfunction via Aldosterone-Dependent Mechanisms in Obese Female Mice. Hypertension (Dallas, Tex 1979). 2016;67(5):1020–8.

50. Rahmouni K, Morgan DA, Morgan GM, Mark AL, Haynes WG. Role of Selective Leptin Resistance in Diet-Induced Obesity Hypertension. Diabetes (New York, NY). 2005;54(7):2012–8.

51. Hall JE, do Carmo JM, da Silva AA, Wang Z, Hall ME. Obesity-Induced Hypertension: Interaction of Neurohumoral and Renal Mechanisms. Circulation research. 2015;116(6):991-1006.

52. Fu Z, Wang F, Liu X, Hu J, Su J, Lu X, et al. Soluble (pro) renin receptor induces endothelial dysfunction and hypertension in mice with diet-induced obesity via activation of angiotensin II type 1 receptor. Clinical Science. 2021;135(6):793–810.

53. Gatineau CE, Gong CM, Yiannikouris CF. Soluble Prorenin Receptor Increases Blood Pressure in High Fat-Fed Male Mice. Hypertension. 2019;74(4):1014–20.

54. Visniauskas B, Reverte V, Abshire CM, Ogola BO, Rosales CB, Galeas-Pena M, et al. High-plasma soluble prorenin receptor is associated with vascular damage in male, but not female, mice fed a high-fat diet. American journal of physiology Heart and circulatory physiology. 2023;324(6):H762–H75.

