## Supplementary data for "Renal-Derived Human sPRR Does Not Increase Blood Pressure in High Fat Diet Mice"

**Supplementary material**

**Tables**

Table 1. Physiological parameters in male and female mice on high fat diet

|  | Male CTL | Male CD-HsPRR | Female CTL | Female CD-HsPRR |
| --- | --- | --- | --- | --- |
| Bodyweight (g) | 47.0±1.5 | 46.2±3.7 | 34.2±2.5 | 40.0±2.2 |
| Kidney weight (% of BW) | 1.0 ±0.04 | 1.1±0.07 | 1.0±0.10 | 1.0±0.04 |

Data are mean ± SEM of 6-8 mice/group; A two-way ANOVA was performed to detect differences. * P<0.05 compared with CTL.

Table 2. Blood pressure parameters (24 h) in male and female mice on high fat diet

|  | Male CTL | Male HsPRR | Female CTL | Female HsPRR |
| --- | --- | --- | --- | --- |
| SBP (mmHg) | 137±1.5 | 138±1.8 | 133±1.5 | 134±1.1 |
| DBP (mmHg) | 102±3.1 | 102±0.9 | 98±1.3 | 98±1.7 |
| MAP (mmHg) | 120±2.0 | 121±1.3 | 116±1.1 | 116±1.1 |
| HR (bpm) | 597±9.3 | 592±15.2 | 645±5.7 | 623±5.8 |

Data are mean ± SEM of 6-8 mice/group; A two-way ANOVA was performed to detect differences. * P<0.05 compared with CTL.
